# Whole genome sequencing of three mesorhizobia isolated from northern Canada to identify genomic adaptations promoting nodulation in cold climates

**DOI:** 10.1101/2022.04.26.489235

**Authors:** Yi Fan Duan, Paul Grogan, Virginia K Walker, George C diCenzo

## Abstract

The N2-fixing symbiosis between rhizobia and legumes is negatively impacted by numerous stresses, including low temperatures. To identify genomic features and biochemical pathways of rhizobia that could foster improved symbiotic function under low temperatures, we isolated and characterized three *Mesorhizobium* strains from legume nodules collected at two distant northern Canadian sites. Whereas the classical determinants of nodulation and nitrogen fixation are located on the chromosome of most mesorhizobia, whole genome sequencing revealed that these genes are on a large symbiotic megaplasmid in all three of the newly isolated strains. A pangenome-wide association study identified 25 genes putatively associated with mesorhizobia isolated from arctic or subarctic environments, with the genomic location of many of these genes implying a relationship with legume symbiosis. Phylogenetic and sequence analyses of the common nodulation genes revealed alleles that are highly conserved amongst mesorhizobia isolated from northern climates but uncommon in mesorhizobia isolated from similar plant hosts in other climatic regions, suggesting potential functional adaptive differences and the horizontal transfer of these alleles between northern rhizobia. We speculate that *nod* sequence divergence was driven by climatic factors, and that the encoded proteins may be particularly stable and/or active at low temperatures.

## INTRODUCTION

The rhizobia are a polyphyletic group of soil-dwelling *α*-proteobacteria and *β*-proteobacteria capable of entering N2-fixing endosymbiotic relationships with leguminous plants. During the symbiosis, the rhizobia are located within a specialized organ known as a root nodule, where they convert atmospheric N2 gas into ammonia for use by the plant as a bio-available source of nitrogen. This process of symbiotic nitrogen fixation (SNF) has the potential to fulfill the complete nitrogen requirement of a plant and is thus a cornerstone of sustainable agricultural practices. For example, it was estimated that crop, pasture, and fodder legumes collectively fix 33 to 46 Tg of nitrogen per year (Herridge et al. 2008). However, establishment of the symbiotic relationship is an intricate process involving multiple steps, each of which can be negatively impacted by adverse environmental conditions (Valentine et al. 2018). Successful inoculation of crops therefore necessitates the identification of rhizobia adapted to local agroclimatic conditions that can overcome the detrimental impacts of relevant stresses.

One environmental stress known to be detrimental to SNF is low temperature (Jones and Tisdale 1921; Prévost et al. 1999). Low soil temperatures result in the formation of fewer nodules and reduced nodule mass (Jones and Tisdale 1921; Rice et al. 1995). This occurs at least in part by interfering with the synthesis of key signal compounds required to establish the symbiosis; low temperatures both reduce flavonoid production by legumes and decrease Nod factor biosynthesis by rhizobia (McKay and Djordjevic 1993; Zhang and Smith 1996a; Zhang et al. 1996). Low temperatures also impair the rate of nodule development, resulting in reduced rates of nitrogen fixation even when normalized by nodule mass (Fyson and Sprent 1982; Prévost et al. 1987a). Although low temperatures also result in a slight decrease in the rate of infection thread growth, this appears to be inconsequential (Fyson and Sprent 1982). In addition to their direct effect on SNF, low temperatures may also indirectly impact SNF, for example, by reducing the bioavailability of soil phosphorus (Hernández et al. 2009; Shaw and Cleveland 2020).

The effects of low temperatures on SNF are particularly relevant in temperate climatic regions, where the suboptimal soil temperatures of spring can result in delayed nodulation that has lasting impacts on the total amount of nitrogen fixed throughout the growing period (Lynch and Smith 1993, 1994; Zhang et al. 1995). The negative impacts of low temperatures can be partially overcome through: i) the addition of exogenous flavonoids (Zhang and Smith 1995, 1996b); ii) the use of rhizobium inoculants that constitutively produce Nod factors (Zhang et al. 2002); or iii) the identification and application of naturally cold-adapted rhizobia (Prévost et al. 1999, 2003). For example, clover rhizobia isolated from more northern regions of Nordic countries displayed more efficient symbiotic properties than clover rhizobia isolated from more southern regions, but only at low temperatures (Ek-Jandér and Fåhraeus 1971; Lipsanen and Lindström 1986). Similarly, sainfoin (*Onobrychis viciifolia*) inoculated with rhizobia isolated from the Arctic displayed higher rates of nodulation and N_2_-fixation at low temperatures compared to sainfoin inoculated with temperate rhizobia (Prévost et al. 1987a; Prévost and Bromfield 1991). Notably, two cold-adapted strains of *Bradyrhizobium japonicum* outperformed the commercial inoculant *B. japonicum* 532 C (originally isolated from soybean in Brazil following inoculation with strains from the United States of America) in field trials performed in Québec, Canada (Zhang et al. 2003).

Understanding the molecular mechanisms for improved SNF under low temperatures could enable the engineering of low temperature tolerance into highly effective rhizobium inoculants. However, this has been hampered by the limited availability of whole genome sequences for rhizobia isolated from cold environments. Here, we report the isolation of three *Mesorhizobium* strains from the nodules of native legumes growing in two distant Low Arctic sites in northern Canada. Whole genome sequencing followed by phylogenetics and comparative genomics was performed to identify genomic adaptations potentially associated with promoting successful nodulation and nitrogen fixation at low temperatures.

## MATERIALS AND METHODS

### Bacterial strains and growth conditions

Bacterial strains used in this work are listed in **Table 1**. All strains were routinely grown using TY medium (5 g L^-1^ tryptone, 2.5 g L^-1^ yeast extract, 10 mM CaCl_2_) or YEM medium (10 g L^-1^ mannitol, 1 g L^-1^ yeast extract, 1.71 mM NaCl, 2.87 mM K_2_HPO_4_, 0.81 mM MgSO_4_). The MM9-mannitol minimal medium consisted of 40 mM 3-(*N*-morpholino)propanesulfonic acid, 20 mM KOH, 19.2 mM NH_4_Cl, 8.76 mM NaCl, 2 mM KH_2_PO_4_, 1 mM MgSO_4_, 0.25 mM CaCl_2_, 38 µM FeCl_3_, 42 nM CoCl_2_, 1 µg/mL biotin, and 10 mM mannitol. The carbon-free minimal medium used for the Phenotype MicroArray analyses was the same as MM9-mannitol except that mannitol was excluded. Unless stated otherwise, strains were grown at 27.5°C.

**Table 1.** Bacterial strains used in this study.

| Organism | Host plant * | Isolation site | Isolation year | Reference |
| --- | --- | --- | --- | --- |
| *Mesorhizobium* sp. AR02 | *Oxytropis nigrescens* | NWT, Canada | 2019 | This study |
| *Mesorhizobium* sp. AR07 | *Oxytropis nigrescens* | NWT, Canada | 2019 | This study |
| *Mesorhizobium* sp. AR10 | Likely *Astragalus* sp. † | YT, Canada | 2019 | This study |
| *Mesorhizobium japonicum* MAFF303099 | *Lotus japonicum* | Tochigi Prefecture, Japan | 1981 | Sawada, Y. Unpublished |

\* Host plant was determined based on morphology, with genus designations confirmed via Sanger sequencing of the *matK* gene.
† Please refer to the Results and Discussion subsection “Isolation of mesorhizobia from northern Canada” for a discussion of the uncertainty around the host plant of *M. sp.* AR10.

### Isolation of rhizobia

Whole *Oxytropis nigrescens* plants including root systems were collected at two similar esker-top (N 64° 51’ 22.7”; W 111° 39’ 23.0”) locations (∼ 100 m apart) near the Daring Lake Research Station (Northwest Territories, Canada; 64° 52’ N, 111° 33’ W) in mid-August 2019 (photographs of the collection site are provided as **Figure S1A-D**). At these sampling sites, mean diel soil temperature ranges from 0 up to a maximum of 5°C throughout the growing season, with an average annual air temperature of -8°C, and typical annual precipitation between 200 and 300 mm with 138 mm on average falling as rain primarily in June to September (Lafleur and Humphreys 2008; Nobrega and Grogan 2008; Zamin et al. 2014). A second sampling of an assortment of whole plants was performed at the edge of a forest near the Yukon River approximately 25 km north of Whitehorse, Yukon, Canada in October 2019 (a photograph of the collection site is provided as **Figure S1E**). This region is characterized by mean surface soil temperature of 0.5°C, a yearly average air temperature of -0.2°C (with a range of 6-14°C in the growing season), and an annual precipitation of 240 mm, again mostly falling as rain during the summer months (Roy et al. 2021).

Plant material was brought back to Queen’s University (Kingston, Ontario, Canada), where leaf tissue was stored at -80°C until use. Nodules were individually rinsed with double-distilled water (ddH_2_O). Individual, rinsed nodules were surface sterilized with 1% hypochlorite for 15 min, and then rinsed twice with YEM broth. Each nodule was individually crushed in 100 µL of YEM broth within sterile 1.5 mL tubes, following which the media was serial diluted and spread plated on YEM agar plates. Plates were incubated at 27.5°C until colonies formed. Individual colonies were streak purified three times on YEM medium, following which -80°C frozen stocks were prepared.

### Symbiosis assays

Leonard jar assemblies were constructed as described previously (diCenzo et al. 2015). Sainfoin seeds (*Onobrychis viciifolia*; Speare seeds, Harriston, ON, Canada) were dehulled with a KitchenAid Coffee Grinder (product # BCG111OB) and then sterilized with 95% ethanol for five min followed by 2.5% hypochlorite for 20 min. Sterilized seeds were rinsed and soaked with sterile ddH_2_O for 90 min at room temperature. Seeds were germinated for 48 h on 1.5% water agar at room temperature in the dark. Three seedlings were then potted in each Leonard assembly and grown for two nights. Next, *Mesorhizobium* strains grown in TY medium were centrifuged (16,000 *g*, 60 sec, room temperature), and washed and resuspended with sterile 0.85% NaCl to an optical density at 600 nm (OD_600_) of 0.5. One mL of each cell suspension was added to 30 mL of sterile ddH_2_O, and 10 mL added to each Leonard assembly. Plants were grown in an AC-60 chamber (Enconair; now called BioChambers) for 36 days with a day (18 h, 21°C, light intensity of 300 µmol m^-2^ s^-1^) and night (6 h, 17°C) cycle. At the end of the growth period, bacteria were isolated from nodules as described above, and 16S rRNA gene fragments were amplified using polymerase chain reaction (PCR) and sequenced with Sanger sequencing.

### Analysis of growth characteristics

Growth curves in TY broth were performed in a Synergy H1 plate reader (BioTek) largely as described previously (diCenzo et al. 2014). Strains pre-grown in TY broth were diluted to an OD_600_ of ∼ 0.05 in fresh TY broth, and 200 µL added to four replicate wells of a 96-well microtitre plate. The microtitre plate was incubated in the plate reader with shaking at 27.2°C to 27.3°C with a 1°C vertical gradient to limit condensation.

To examine growth dynamics at various temperatures, strains were pre-grown in TY broth at 27.5°C. Strains were diluted to an OD_600_ of 0.05 in 10 mL of fresh TY broth, and 1 mL of each cell suspension was each added to 6 sterile test tubes. Three test tubes were placed in a Thermo Scientific Cel-Gro tissue culture rotator and incubated at 27.5°C. The remaining three test tubes were inserted into a test tube rack on an angle and then incubated with shaking (200 rpm) in a I24R shaker (Eppendorf) for 20 to 48 h at the desired temperature (32.5°C, 30°C, 27.5°C, 25°C, 22.5°C, 20°C, 15°C, or 10°C). At the end of the growth period, 200 µL from each culture was transferred to a 96-well microtitre plate, and the OD_600_ measured using a Synergy H1 plate reader. OD_600_ values were standardized to a 1 cm pathlength through multiplying values by 1.78 as determined from a standard curve.

### Phenotype MicroArrays

Carbon source utilization patterns of *Mesorhizobium* strains were determined using columns one through nine of Gen III plates (Biolog). Cultures pre-grown in MM9-mannitol medium were pelleted by centrifugation (16,000 *g*, 60 sec, room temperature), washed once with carbon-free MM9 medium, and resuspended in carbon-free MM9 to an OD_600_ of ∼ 0.02. Each cell suspension (10 mL) was mixed with 102 µL of Dye Mix A (Biolog), and 100 µL added to each well of a Gen III plate. Plates were taped closed to prevent evaporation, and the absorbance in each well at 590 nm (to monitor reduction of Dye Mix A) and at 750 nm (as reference) were measured using a Synergy H1 plate reader. Plates were incubated without shaking at 27°C, and absorbance at both wavelengths was measured daily for 10 days in a Synergy H1 plate reader. Prior to each measurement, the plates were shaken for five sec in the plate reader. An increase in absorbance at 590 nm was interpreted as indicating the strain could catabolize the corresponding carbon substrate.

### 16S rRNA and *matK* gene sequencing

Bacterial 16S rRNA genes were PCR amplified from individual colonies using the 27f and 1495r universal primers (Lane 1991) using *Taq* FroggaMix (Froggabio), an annealing temperature of 49°C, and an elongation time of 90 sec. Sanger sequencing of the PCR products was performed using the primers 27f and 1495r at the Centre de Recherche, CHU de Québec, Université Laval (Québec City, Québec, Canada).

Approximately 25 mg of plant leaf tissue was ground in 100 µL of T10E1 buffer (10 mM Tris, 1 mM EDTA, pH 8.0) and incubated in a water bath at 60°C for 10 min, as described elsewhere (Ben-Amar et al. 2017). Tubes were briefly centrifuged, and the supernatant transferred to a new tube and stored at -20°C until use. The plant mitochondrial *matK* gene was PCR amplified from those solutions using primers trnK685F and trnk2R (Wojciechowski et al. 2004), Q5 DNA polymerase with the Q5 GC enhancer (New England Biolabs), an annealing temperature of 60°C, and an elongation time of 60 sec. To increase yield, a second round of PCR was performed using 0.5 µL of the original PCRs as the template. Sanger sequencing was performed using the primer matK4La (Wojciechowski et al. 2004) at the Centre de Recherche, CHU de Québec, Université Laval. All primer sequences are provided in **Table S1**.

### Genome sequencing

Two 3 mL TY cultures per strain were inoculated from single colonies and incubated in a rotator drum until stationary phase. Total genomic DNA was isolated from each 3 mL culture using phenol – chloroform – isoamyl alcohol extractions followed by ammonium acetate – isopropanol precipitations as described elsewhere (Cowie et al. 2006), and the RNase A treated DNA pellets were resuspended in T10E1 buffer. Prior to Oxford Nanopore sequencing, an aliquot of DNA from each sample was precipitated and resuspended in commercial nuclease-free water (FroggaBio).

Oxford Nanopore sequencing was performed in-house. Samples were prepared for sequencing using a Rapid Sequencing Kit (SQK-RAD004) following the manufacturer’s instructions. Sequencing was performed using a minION with R9.4.1 flow cells, and the MinKNOW software. Basecalling was performed using GPU-enabled Guppy version 5.011+2b6dbffa5 and the high accuracy model (Oxford Nanopore Technologies). Illumina sequencing (150 bp paired-end reads) was performed at the Microbial Genome Sequencing Center (Pittsburgh, PA, USA) using a NextSeq 550 instrument.

### Genome assembly and annotation

Draft genome assemblies were prepared with Flye version 2.8.3-b1725 (Kolmogorov et al. 2019) and the Nanopore reads. Assembly was also tested using Canu version 2.2-development (r10239) (Koren et al. 2017). Canu and Flye gave similar assembly structures for two of the three strains; however, unlike Flye, Canu failed to fully assembly the genome of *Mesorhizobium* sp. AR10 likely due to the low coverage of ∼ 17.5x. Assemblies generated by Flye were checked for overlaps between contig ends using NUCmer version 4.0.0rc1 (Kurtz et al. 2004), and removed if identified. Genome assemblies were then polished using a multistep process. First, the genome assemblies were polished using Racon version 1.4.22 (Vaser et al. 2017) using Nanopore reads aligned to the draft assemblies with Minimap2 version 2.20-r1061 (Li 2018). The genome assemblies were further polished using Medaka version 1.4.1 (Oxford Nanopore Technologies) and the Nanopore reads. Next, Pilon version 1.24 (Walker et al. 2014) was used to polish the assemblies using the Illumina reads mapped to the assemblies with bwa version 0.7.17-r1198-dirty (Li and Durbin 2009). A final round of polishing was performed using Racon and the Illumina reads, which were aligned to the genome assemblies using bwa. The assemblies were again checked for overlaps between contig ends and removed if found. Finally, replicons were reoriented using Circlator version 1.5.5 (Hunt et al. 2015).

Genome assemblies were annotated using a local copy of the Prokaryotic Genome Annotation Pipeline version 2021-07-01.build5508 (Tatusova et al. 2016). Putative prophage regions were predicted using PhiSpy version 4.2.6 with 1000 random forest trees (Akhter et al. 2012), and putative CRISPR loci were predicted using Prokka 1.14.6 (Seemann 2014).

### Species phylogenetic analyses

The RefSeq version of all 94 *Mesorhizobium* genomes that were either classified as a RefSeq representative genome or that had an assembly level of ‘complete’ or ‘chromosome’ were downloaded from the National Center for Biotechnology Information (NCBI) Genome database. In addition, six complete *Brucella* genomes were downloaded to serve as an outgroup for the rooted phylogeny. All genomes were downloaded on 27 July 2021, and the associated metadata is available as **Datasets S1 and S2**.

A rooted, maximum likelihood phylogeny was constructed from a multilocus sequence analysis using a previously published pipeline (Fagorzi et al. 2020) dependent on AMPHORA2 (Wu and Scott 2012), MAFFT version 7.310 (Katoh and Standley 2013), trimAl version 1.4rev22 (Capella-Gutiérrez et al. 2009), and RAxML version 8.2.12 (Stamatakis 2014). RAxML was run with the GAMMA model of rate heterogeneity and the LG amino acid substitution model. The LG substitution model was chosen as a preliminary run of RAxML with the automatic model selection indicated that it provided the best scoring tree. The final tree represents the bootstrap best tree following 250 bootstrap replicates, with bootstrapping stopped based on the extended majority-rule consensus tree criterion.

To construct an unrooted core-genome gene phylogeny, a pangenome of the *Mesorhizobium* strains was calculated using Roary version 3.12.0 (Page et al. 2015) with a percent identify threshold of 70%. Prior to pangenome calculation, all genomes were reannotated with Prokka to ensure consistent annotation and to prepare files correctly formatted for use with Roary. As part of Roary, the nucleotide sequences of the 646 core genes were individually aligned with MAFFT and the alignments concatenated. The concatenated alignment was trimmed using trimAl, and then used to construct a maximum likelihood phylogeny with RAxML and the GTRCAT model. The final tree represents the bootstrap best tree following 50 bootstrap replicates, with bootstrapping stopped based on the extended majority-rule consensus tree criterion. Phylogenies were visualized with the online iTOL webserver (Letunic and Bork 2016).

Pairwise average nucleotide identity (ANI) values were calculated using FastANI (Jain et al. 2018) with default parameters.

### Pangenome construction and pangenome-wide association studies

The 97 *Mesorhizobium* genomes (the three newly sequenced strains plus the 94 genomes downloaded from NCBI) were reannotated with Prokka prior to pangenome calculation to ensure consistent annotations. Pangenomes were determined using Roary with percent identity thresholds of 90%, 80%, 70%, 60%, 50%, and 40%. Scoary (Brynildsrud et al. 2016) was used to perform pangenome-wide association studies to identify genes associated with rhizobia isolated from regions with arctic or sub-arctic climates; for these analyses, *M*. sp. AR02, *M*. sp. AR07, *M*. sp. AR10, *Mesorhizobium loti* 582, and *Mesorhizobium huakuii* 583 were annotated as having the relevant trait, while the other 92 genomes were annotated as not having the relevant trait. Genes that were annotated by Prokka but that were either not annotated or annotated as pseudogenes by PGAP, were removed from the Scoary output.

### Phylogenetic analyses of symbiotic nitrogen fixation genes

The proteomes were collected for the 97 *Mesorhizobium* strains (the three newly sequenced strains plus the 94 downloaded from NCBI) and combined. Proteomes were searched for NodA, NodB, NodC, NifH, NifD, and NifK proteins using a previously described pipeline (Fagorzi et al. 2020) dependent on the Pfam version 34.0 (Finn et al. 2016) and TIGRFAM version 1.50 (Haft et al. 2013) hidden Markov model databases, as well as HMMER version 3.1b2 (Eddy 2009). Each set of orthologs was aligned using MAFFT, and percent identity matrixes calculated using the phytools package of R (Revell 2011).

The aligned NodA, NodB, and NodC orthologs were trimmed with trimAl, and then used to construct maximum likelihood phylogenies with RAxML. For NodA, the JTT amino acid substitution model was used, and 400 bootstrap replicates performed. For NodB, the DUMMY2 amino acid substitution model was used, and 512 bootstrap replicates performed. For NodC, the DUMMY2 amino acid substitution model was used, and 352 bootstrap replicates performed. In all cases, the GAMMA model of rate heterogeneity was used, and the final tree represents the bootstrap best tree. The amino acid substitution models were chosen based on the results of preliminary runs of RAxML with the automatic model selection option, while the number of bootstrap replicates was determined by the extended majority-rule consensus tree criterion.

### Sequence-based clustering of the common nodulation genes

The nucleotide sequence of all NCBI Nucleotide bacterial entries containing either *nodA* (4,828 entries), *nodB* (676 entries), or *nodC* (7,824) in the title were downloaded on 30 July 2021, regardless of whether they were complete or partial sequences (**Datasets S3-S5**). The datasets were supplemented with the *nodA, nodB*, and *nodC* genes of *M.* spp. AR02, AR07, and AR10. Each set of orthologs was clustered using CD-HIT version 4.8.1 (Li and Godzik 2006) with a percent identity threshold of 95% and at least 80% coverage of the smaller gene.

## RESULTS AND DISCUSSION

### Isolation of mesorhizobia from northern Canada

Native *O. nigrescens* plants growing in the Low Arctic near Daring Lake, located 300 km northeast of Yellowknife, NWT, Canada, and from plants growing in a taiga forest close to the Yukon River, 25 km north of Whitehorse, YT, Canada, were examined for the presence of root nodules. These tundra and taiga sites, respectively, are classified as plant hardiness zone 0 (temperatures as low as -54°C). Nine *O. nigrescens* plants at the NWT site were uprooted, five of which had at least one structure resembling a nodule. Rhizobia were successfully isolated from two of these putative nodules, and given the strain names AR02 and AR07 (**Table 1**). Only one pink nodule was identified after examining the roots of several plants at the YT site site; the rhizobium isolated from this nodule was named AR10 (**Table 1**). The YT sample contained more than one vascular plant, including *Lupinus arcticus*, but subesquent PCR amplification and Sanger sequencing of the *matK* gene indicated the presence of an *Astragalus* sp. Given that AR10 was able to nodulate sainfoin and carries nod genes highly similar to other *Astragalus* and *Oxytropis* micosymbionts (discussed below), and that the nodule had a classical indeterminate morphology as opposed to a lupinoid morpholophy, we therefore attribute strain AR10 to an *Astragalus* plant of an unidentified species. Sequencing of the 16S rRNA gene indicated that all three newly-isolated rhizobia belong to the genus *Mesorhizobium*. In addition, all three strains were able to nodulate sainfoin (*O. viciifolia*) (**Figure S2**), consistent with other mesorhizobia isolated from *Oxytropis* and *Astragalus* plants in northern Canada (Prévost et al. 1987b).

### Growth characteristics of *M.* sp AR02, AR07, and AR10

Large variation was observed in the growth rates of *M.* spp. AR02, AR07, and AR10 in TY medium at 27°C. *M.* sp. AR02 was the fastest growing strain with a generation time of 5.3 ± 0.1 h (average ± standard deviation), which was slightly slower than the generation time of the common laboratory strain *Mesorhizobium japonicum* MAFF303099 isolated in Japan (5.0 ± 0.1 h). *M.* sp. AR07 and AR10 had generation times of 10.3 ± 0.2 and 6.1 ± 0.1 h, respectively. As growth rates were not measured on other media, we cannot say whether *M.* sp. AR10 is inheritely a slow growing strain or whether other media would support a growth rate more similar to that of *M.* spp. AR02 and AR10.

Given the geographical isolation sites of *M.* sp. AR02, AR07, and AR10, we hypothesized that their permissible growth temperature ranges would be lower than that of *M. japonicum* MAFF303099, which was isolated from a humid sub-tropical climatic region. Indeed, unlike *M. japonicum* MAFF303099, the three low arctic strains were unable to grow at 32.5°C, and the growth of *M.* sp. AR10 was also inhibited at 30°C (**Figure 1**). The optimal growth temperature for *M.* spp. AR02 and AR07 was 30°C, while the optimal growth temperature for *M.* sp. AR10 was 27.5°C (**Figure 1**). *M.* sp. AR10, which was the most sensitive strain to high temperatures, appeared to be the fastest growing of the three newly isolated strains at 15°C and 10°C (**Figures 1, S3**). *M. japonicum* MAFF303099 and *M.* sp. AR02 appeared most sensitive to low temperatures while growth of *M.* sp. AR07, which was the slowest growing strain, appeared to be the least impacted by temperature (**Figures 1, S3**); the doubling times of *M. japonicum* MAFF303099, and *M.* spp. AR02, AR07, and AR10 at 10°C were 8.7, 8.3, 5.5, and 6.6-fold lower than that at 27.5°C, respectively. The ability of all three northern strains as well as MAFF303099 to grow at 10°C means all four strains are classified as psychrotolerant. Overall, these results suggest that *M*. spp. AR02, AR07, and AR10 do not have obvious superior growth characteristics at low temperatures compared to some other mesorhizobia. However, we cannot rule out that differences would emerge at temperatures closer to 0°C, or if they would show higher infection, higher nodulation frequency or maturation, or even more nitrogenase activity compared to standard strains at lower temperatures. It also remains possible that *M*. spp. AR02, AR07, and AR10 may be more capable of long-term survival at freezing temperatures compared to other strains.

**Figure 1.**
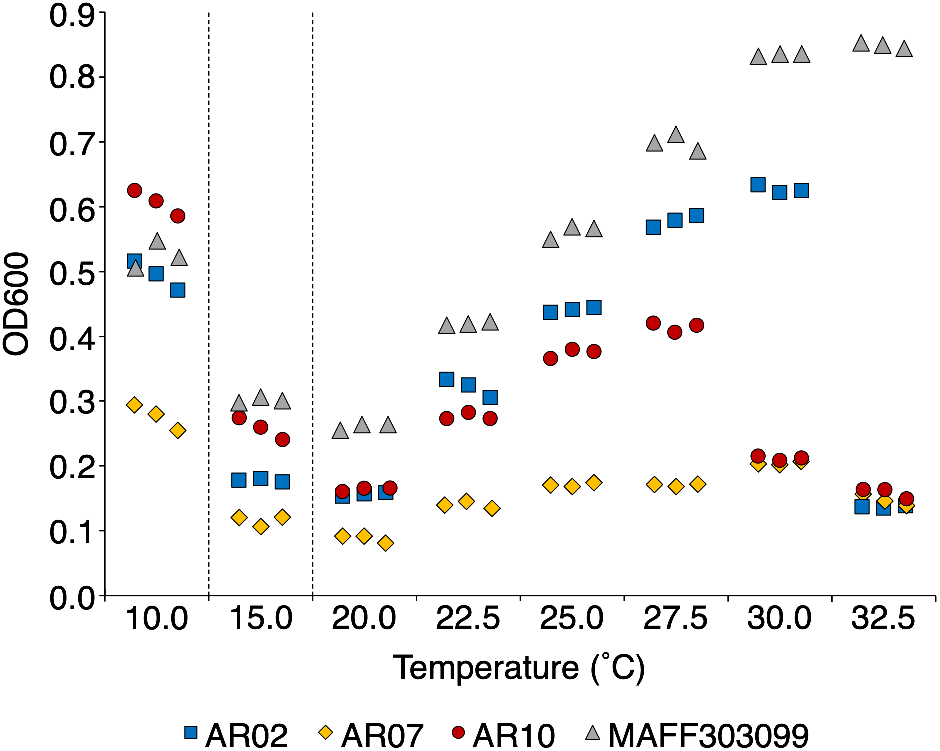
Impact of temperature on growth of *Mesorhizobium* strains. Cell density (OD_600_) of *Mesorhizobium* cultures following ∼ 7 days (10°C), ∼ 2 days (15°C), or ∼ 1 day (20°C to 32.5°C) of growth at the indicated temperature. Starting OD_600_ values were ∼ 0.05 for all cultures. Three biological replicates were performed, with each symbol representing a single replicate. Blue squares – *M.* sp. AR02; yellow diamonds – *M.* sp. AR07; red circles – *M.* sp. AR10; grey triangles – *M. japonicum* MAFF303099.

The carbon metabolic capacities of *M.* spp. AR02, AR07, and AR10 were investigated using Biolog Gen III Phenotype MicroArray plates. Each strain appeared capable of catabolizing 30 (AR02) or 36 (AR07, AR10) carbon sources, with 21 substrates catabolized by all three strains. Overall, the highest rates of cellular respiration (as measured by redox dye reduction) were generally observed in wells containing simple sugars or sugar alcohols, suggesting that these are the preferred carbon sources for all three strains. Of the common set of 21 substrates, 18 were carbohydrates – seven monosaccharides (D-glucose, D-fructose, D-fucose, D-galactose, D-mannose, L-fucose, L-rhamnose), four dissaccharides (D-maltose, D-trehalose, D-turanose, and sucrose), five sugar alcohols (D-arabitol, D-mannitol, D-sorbitol, glycerol, myo-Inositol), and two amino sugars (N-acetyl-D-galactosamine, N-acetyl-D-glucosamine) – and three were organic acids (L-malic acid, L-glutamic acid, L-lactic acid). A complete list of carbon substrate catabolized by *M.* spp. AR02, AR07, and AR10 is provided as **Table S2**, and all absorbance measurements are provided as **Datasets S6-S8**.

**Table 2.**
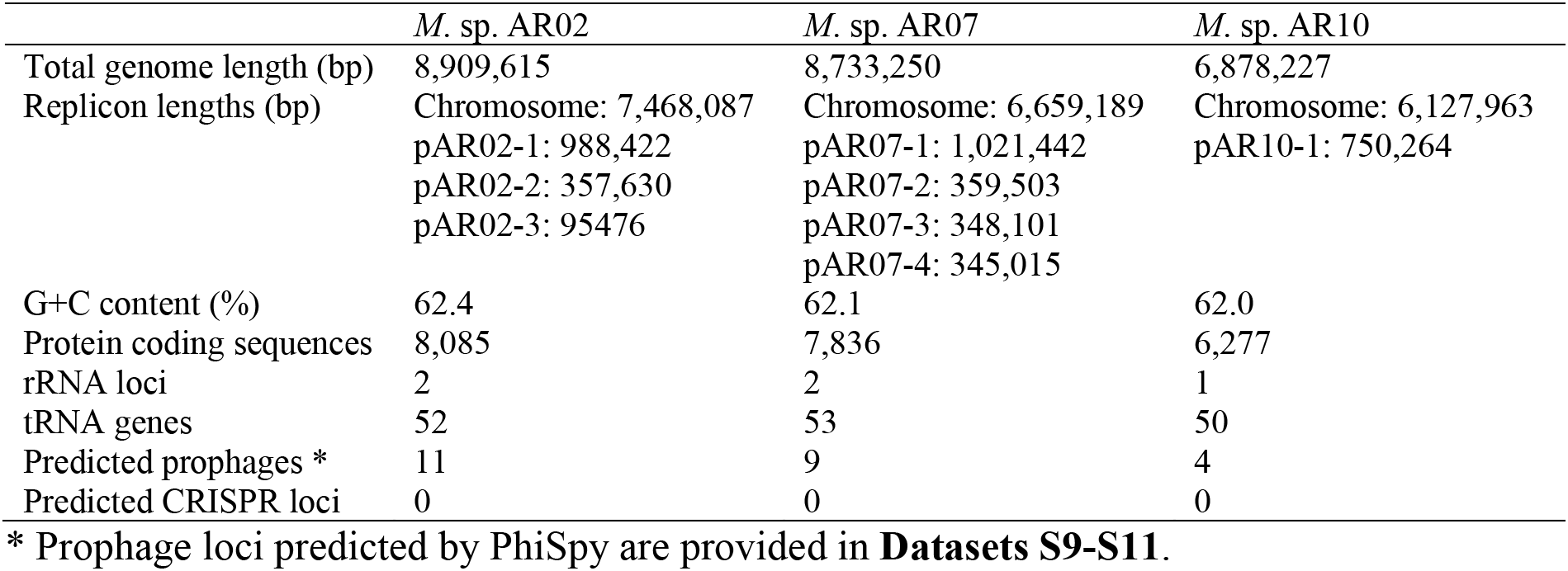
Features of the *M.* sp. AR02, *M.* sp. AR07, and *M.* sp. AR10 genome assemblies generated in this study.

|  | *M. sp.* AR02 | *M. sp.* AR07 | *M. sp.* AR10 |
| --- | --- | --- | --- |
| Total genome length (bp) | 8,909,615 | 8,733,250 | 6,878,227 |
| Replicon lengths (bp) | Chromosome: 7,468,087<br>pAR02-1: 988,422<br>pAR02-2: 357,630<br>pAR02-3: 95476 | Chromosome: 6,659,189<br>pAR07-1: 1,021,442<br>pAR07-2: 359,503<br>pAR07-3: 348,101<br>pAR07-4: 345,015 | Chromosome: 6,127,963<br>pAR10-1: 750,264 |
| G+C content (%) | 62.4 | 62.1 | 62.0 |
| Protein coding sequences | 8,085 | 7,836 | 6,277 |
| rRNA loci | 2 | 2 | 1 |
| tRNA genes | 52 | 53 | 50 |
| Predicted prophages * | 11 | 9 | 4 |
| Predicted CRISPR loci | 0 | 0 | 0 |

\* Prophage loci predicted by PhiSpy are provided in **Datasets S9-S11**.

### Taxonomy and genomic properties of *M.* sp. AR02, AR07, and AR10

To gain insight into the genetic determinants facilitating nodulation at low temperatures, the genomes of *M.* spp. AR02, AR07, and AR10 were sequenced. Closed genomes were obtained for all three strains, and summary statistics are presented in **Table 2**. Maximum likelihood phylogenies constructed from the concatenated alignments of 30 highly conserved proteins (**Figure S4**) or on 646 core genes (**Figure 2**) revealed that *M.* spp. AR02, AR07, and AR10 are not monophyletic but are spread across the genus *Mesorhizobium*. ANI calculations suggest that each strain represent novel genospecies as all comparisons between these strains and the 94 *Mesorhizobium* genomes downloaded from NCBI gave values < 95%.

**Figure 2.**
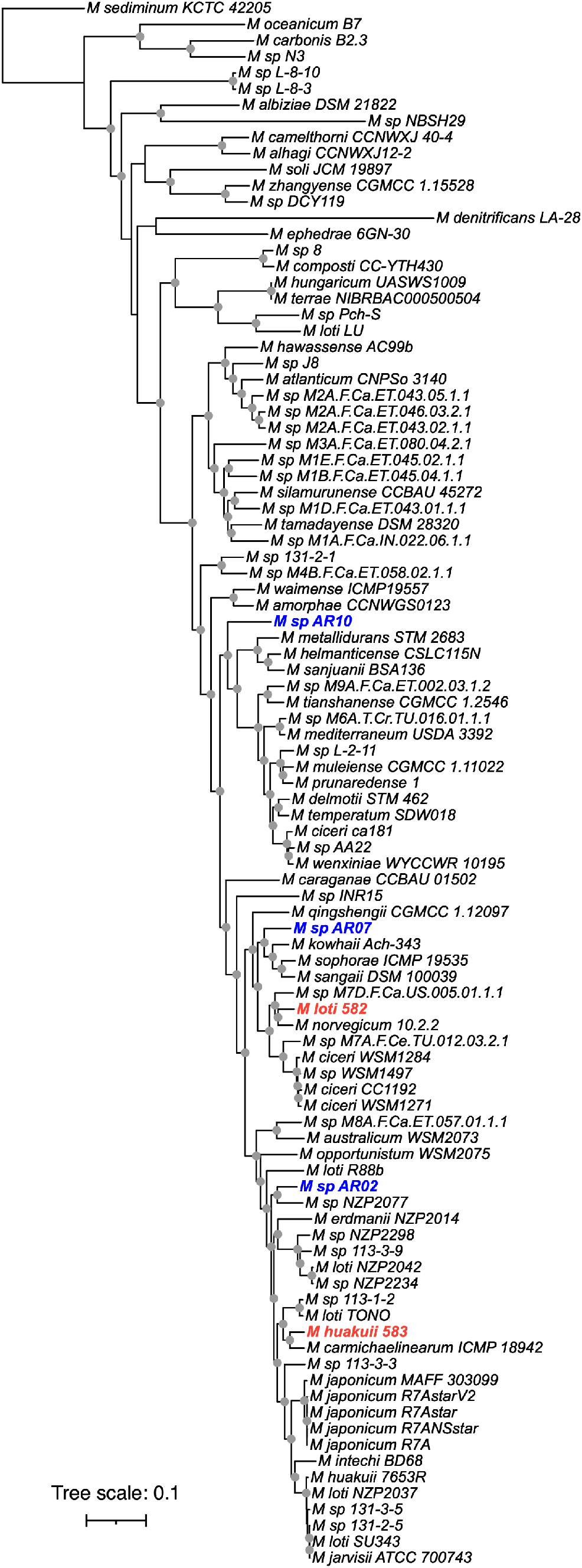
Unrooted phylogeny of the genus *Mesorhizobium.* A maximum likelihood phylogeny of 97 strains was prepared from a concatenated alignment of 646 core genes. Nodes with a bootstrap value of ≥ 90 are indicated with gray dots, based on 50 bootstrap replicates. The scale represents the mean number of nucleotide substitutions per site. Strains isolated and sequenced in this study are shown in blue, boldface font. *M. loti* 582 and *M. huakuii* 583, also isolated from *Oxytropis* plants in a subarctic climatic region, are shown in red, boldface font.

The genome structures of the two *Oxytropis* isolates (AR02, AR07) collected near Daring Lake are unusually large and divided for members of the genus *Mesorhizobium* (**Table 2**). Their genome sizes of 8.9 Mb (8,085 protein coding genes) and 8.7 Mb (7,836 protein coding genes), respectively, are nearly 2 Mb larger than the average *Mesorhizobium* genome (6.9 Mb carrying 6,361 protein coding genes, based on 61 fully assembled genomes available through NCBI) and exceed the size of the next largest fully assembled genome in the NCBI database (*M. loti* TONO; 8.5 Mb carrying 7,831 protein coding genes). Additionally, *M.* sp. AR02 and AR07 carry three and four plasmids, respectively; in comparison, only ∼ 23% of the other 61 finished *Mesorhizobium* genomes have more than one plasmid, while only ∼ 11% have three or more plasmids. Finally, we examined the G+C content of *M.* spp. AR02, AR07, and AR10 as we hypothesized that the G+C content of these strains would be lower than average for the mesorhizobia if they have adapted to cold climates. Although the G+C content of the three strains is slightly below average for mesorhizobia (**Table 2**), the values fall within the standard range for the genus and are thus not unusually low (average of 62.6% with a standard deviation of 1.2% for the 1,080 *Mesorhizobium* with finished or draft genomes available through NCBI).

### Symbiotic gene organization

*M*. spp. AR02, AR07, and AR10 carry the nitrogenase genes (*nifH, nifD, nifK*) and the common nodulation genes (*nodA, nodB, nodC*) on large megaplasmids between ∼ 750 kb and ∼ 1 Mb in size. This contrasts with the majority of mesorhizobia that more often carry these genes within a symbiotic island on the chromosome; of the 50 *Mesorhizobium* genomes downloaded from NCBI carrying *nifH, nifD, nifK, nodA, nodB*, and *nodC*, only five had these genes on a plasmid. Interestingly, the only other strains in our dataset isolated from *Oxytropis* (*M. loti* 582 and *M. huakuii* 583, isolated from the Kamchatka Peninsula, Russia) or *Astragalus* plants (*M*. *huakuii* 7653R, isolated from Nanjing, China) also contained these six genes on a megaplasmid (Wang et al. 2014; Safronova et al. 2020). Thus, although symbiotic megaplasmids are uncommon in the genus *Mesorhizobium*, they appear to be the dominant source of the symbiotic genes in microsymbionts of the subtribe Astragalinae.

In all three strains, the common nodulation genes *nodA, nodB*, and *nodC* were found as part of a larger *nod* locus that also contains the nodulation genes *nodF, nodE, nodG, nodI, nodJ, nodH, nodP*, and *nodQ*, as well as a gene of unknown function (DUF2061) (**Figure 3**). The same organization is also found in three other Astragalinae microsymbionts isolated from sub-arctic or arctic climatic regions: *Mesorhizobium* sp. N33 isolated from *Oxytropis arctobia* near Sarcpa Lake (Nunavut, Canada) (Prévost et al. 1987b; Cloutier 1996) and *M. loti* 582 and *M. huakuii* 583 isolated from *Oxytropis kamtschatica* in the Kamchatka Peninsula (Russia) (Safronova et al. 2020). In addition to conservation of gene order, all six strains shared > 96% nucleotide identity across this *nod* locus, indicative of recent horizontal gene transfer of the locus, as is common for symbiotic genes in rhizobia (Sullivan et al. 1995; Pérez Carrascal et al. 2016). Aside from these six strains, we did not identify other rhizobia with the same *nod* locus organization. The most similar organization was that of *M. huakuii* 7653R (**Figure 3**), which is also an *Astragalus* microsymbiont but isolated from a plant growing in the humid subtropical climate of Nanjing, China (Zhang et al. 2000). Despite the similarities in *nod* gene organization and host plant, there was significant sequence divergence (< 76% nucleotide identity) when comparing this *nod* locus of *M. huakuii* 7653R with the corresponding loci of the northern rhizobia (*M.* sp. AR02, *M.* sp. AR07, *M.* sp. AR10, *M*. sp. N33, *M. loti* 582, and *M. huakuii* 583).

**Figure 3.**
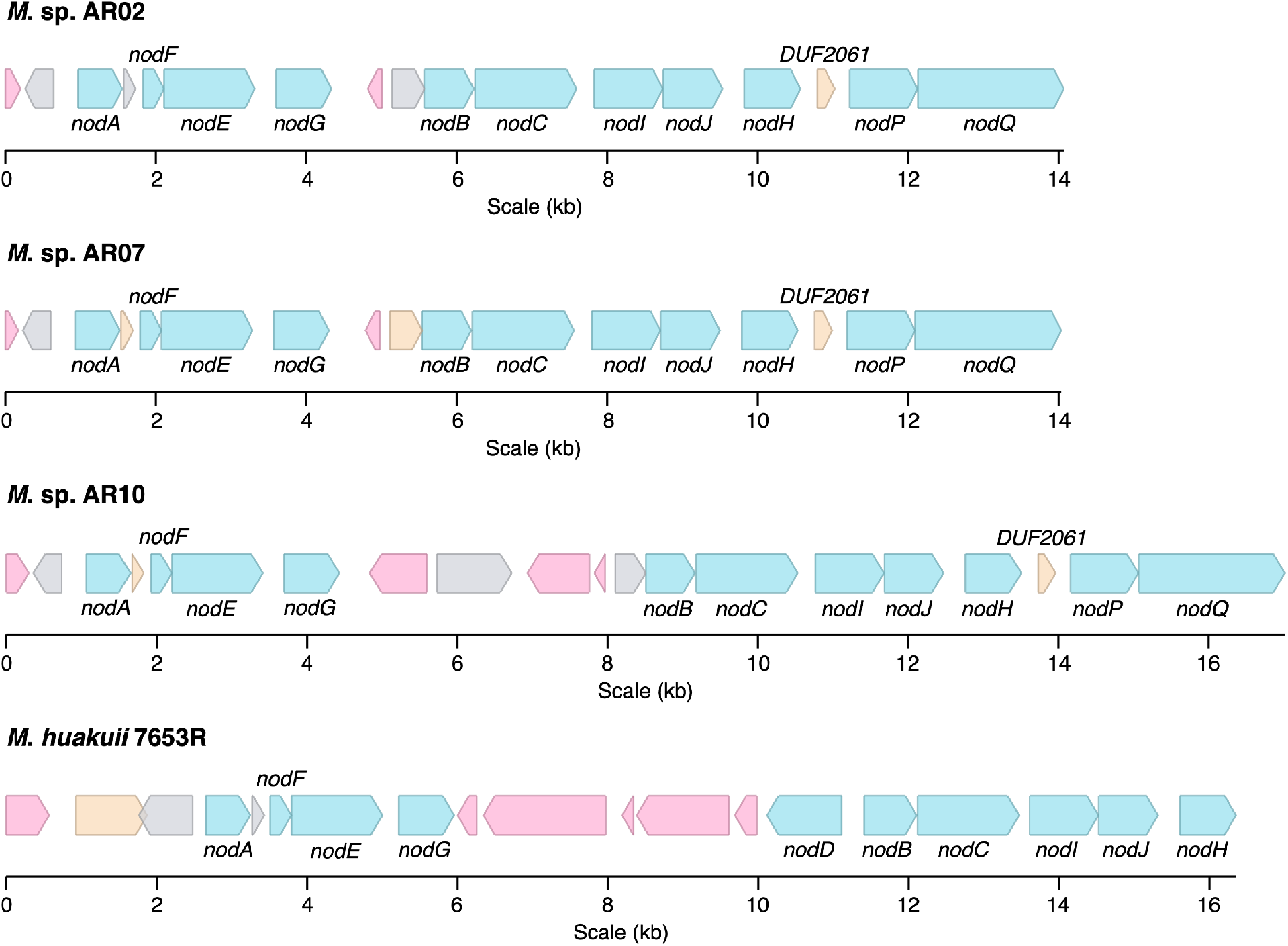
Organization of the *nod* locus encompassing the common nodulation genes of four *Mesorhizobium* strains. The loci encompassing the common nodulation genes (*nodA*, *nodB*, *nodC*) of the three strains isolated and sequenced in this study (*M*. sp. AR02, *M*. sp. AR07, *M*., sp. AR10) and an *Astragalus* microsymbiont isolated from a humid subtropical climate (*M. huakuii* 7653R) are shown. Nodulation genes are shown in blue. Transposases, including pseudogenes showing similarity to transposases, are shown in red. Pseudogenes are shown in grey. All other genes are shown in orange. The pseudogenes downstream of *nodA* in *M*. sp. AR02 and *M. huakuii* 7653R, as well as the pseudogenes upstream of *nodB* in *M.* sp. AR02 and *M.* sp. AR10, were not annotated by PGAP but added to the figure based on sequence conservation with the corresponding hypothetical genes of *M.* sp. AR07. Diagram is drawn to scale.

### Phylogenetics of the common nodulation proteins

The low sequence conservation between the *nod* loci of *M.* spp. AR02, AR07, and AR10 with the corresponding region of *M. huakuii* 7653R prompted us to examine the phylogenetic relationship of the NodA (**Figure S5**), NodB (**Figure S6**), and NodC (**Figure 4**) proteins in our dataset of 97 *Mesorhizobium* genomes (see Materials and Methods). For all three protein families, the proteins from *M.* sp. AR02, *M.* sp. AR07, *M.* sp. AR10, *M. loti* 582, and *M. huakuii* 583 formed a tight cluster with > 97% amino acid sequence identity, while the next most closely related proteins had < 83 % amino acid identity except for one NodC protein with ∼ 87% identity. This result suggested that the *nod* genes of *Mesorhizobium* microsymbionts of Astragaline plants in sub-arctic and arctic climates are highly conserved amongst each other likely as a result of horizontal transfer, but distinct from those in other climates. We hypothesized the prevalence of these *nod* alleles in northern environments was climate-related as plant host range appeared insufficient to explain this result given that the *M. huakuii* 7653R proteins do not group with those of the northern strains (**Figures 4, S5, S6**).

**Figure 4.**
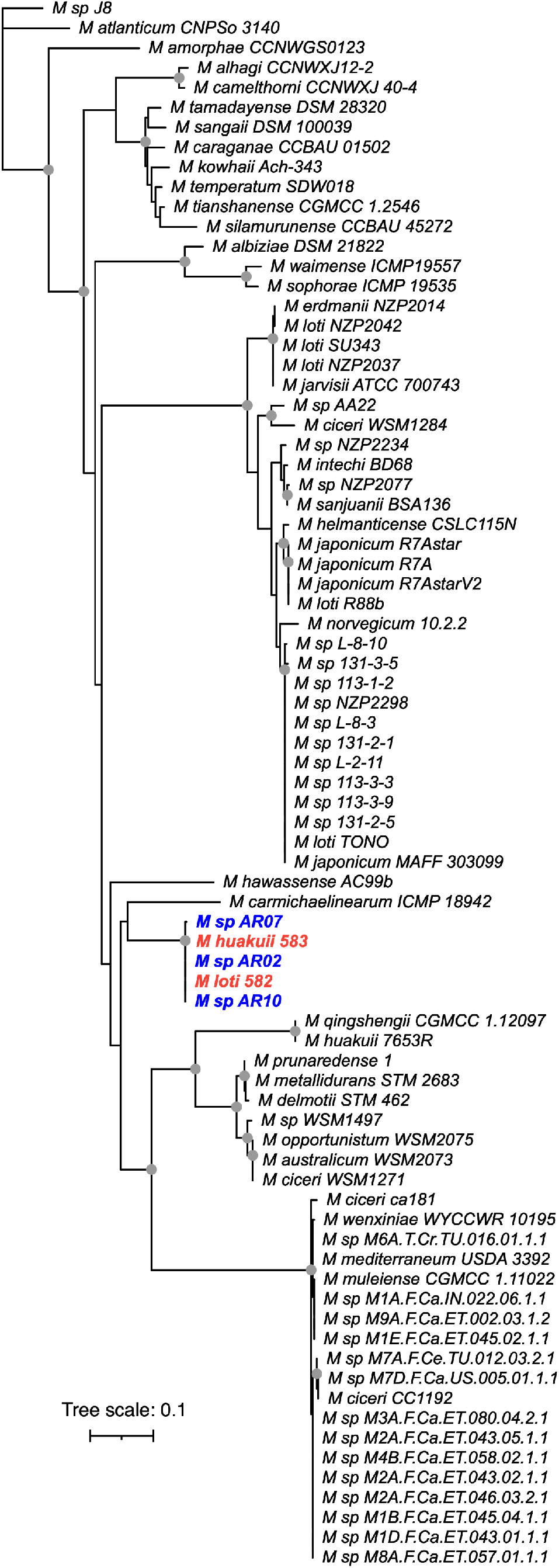
Phylogeny of the NodC proteins of the mesorhizobia. A maximum likelihood phylogeny of the NodC proteins of the 97 *Mesorhizobium* strains shown in Figure 2. Nodes with a bootstrap value of ≥ 90 are indicated with gray dots, based on 352 bootstrap replicates. The scale represents the mean number of amino acid substitutions per site. Strains isolated and sequenced in this study are shown in blue, boldface font. *M. loti* 582 and *M. huakuii* 583, also isolated from *Oxytropis* plants in a subarctic climatic region, are shown in red, boldface font.

To test the above hypothesis, we downloaded the complete or partial nucleotide sequences of 4,837 *nodA* genes, 745 *nodB* genes, and 7,871 *nodC* genes, supplemented the datasets with the corresponding genes of *M*. spp. AR02, AR07, and AR10, and clustered each set of orthologs using a nucleotide identity threshold of 95%. For *nodA*, the genes from *M.* spp. AR02, AR07, and AR10 formed a four-gene cluster with the *nodA* gene of *M.* sp. N33 (**Table S3**) (Cloutier 1996). In the case of *nodB*, the genes from our three strains formed a cluster with 11 additional genes (**Table S3**). These 11 genes included *nodB* from *M.* sp. N33, the *nodB* genes of six rhizobia isolated from root nodules of *Astragalus*, *Oxytropis*, or the related plant genus *Hedysarum* grown in Alaska (USA; subarctic climate) (Wernegreen and Riley 1999), as well as the *nodB* genes of four of five rhizobia isolated from root nodules of *Astragalus* or *Oxytropis* plants grown in Altai or the Moscow region of Russia (humid continental climate) (Wernegreen and Riley 1999). In terms of *nodC*, the genes from *M.* spp. AR02, AR07, and AR10 clustered with an additional 20 genes (**Table S3**). These included the *nodC* gene from the same 11 strains included in the *nodB* cluster, the *nodC* genes of eight of 13 mesorhizobia isolated from root nodules of *Astragalus* or *Oxytropis* plants grown in arctic and sub-arctic climatic regions of Sweden and Russia (Ampomah et al. 2017), and two of 39 mesorhizobia with sequenced *nodC* genes isolated from root nodules of *Astragalus*, *Oxytropis*, or *Hedysarum* plants grown in cold semi-arid or humid continental climatic regions of China (Yan et al. 2016).

Overall, the above results support our hypothesis that the nodulation genes of *M.* spp. AR02, AR07, and AR10 are highly conserved across *Mesorhizobium* microsymbionts of Astragaline (and *Hedysarum*) plants in arctic and sub-arctic climates and may represent the dominant set of *nod* alleles in those climates. This is presumably due in part to the common plant host of these microsymbionts (Roche et al. 1996). However, as noted above, host range seems insufficient to explain the apparent pervasive spread of these *nod* alleles in arctic and subarctic climates as they appear to be rare in microsymbionts of the same hosts in other environments. For example, similar *nodC* alleles were not present in 37 of 39 mesorhizobia isolated from root nodules of *Astragalus*, *Oxytropis*, or *Hedysarum* plants grown in cold semi-arid or humid continental climatic regions of China (Yan et al. 2016). Likewise, i) the *nodC* alleles of 32 of 32 rhizobia (of which 14 were mesorhizobia) isolated from *Astragalus* plants in temperate climatic regions of China (Zhao et al. 2008), ii) all six mesorhizobia isolated from nodules of *Astragalus* plants grown in the Lublin region of Poland (hemiboreal climate) (Gnat et al. 2014, 2015), and iii) the four rhizobia isolated from root nodules of *Astragalus* or *Oxytropis* plants grown in North Dakota (USA; humid continental climate), South Dakota (USA; humid continental climate), Afghanistan (dry climate), and Israel (Mediterranean climate) (Wernegreen and Riley 1999) clustered separately from the *nodC* genes of our mesorhizobia. Thus, we conclude that the *nod* genes of most *Astragalus*, *Oxytropis*, and *Hedysarum* microsymbionts in arctic and subarctic climates are divergent from mesorhizobia in other climatic regions. Although further study is required to determine if the sequence divergence has meaningful impacts on the function of the encoded proteins, it is tempting to speculate that the Nod proteins of the arctic and subarctic mesorhizobia are more active at low temperatures compared to other Nod protein variants, thereby promoting higher rates of nodulation and/or infection, and ultimately nitrogen fixation.

### Genes associated with mesorhizobia isolated from cold climates

To identify genes putatively associated with growth, nodulation, and/or nitrogen fixation at low temperatures, a pangenome of 97 *Mesorhizobium* strains was calculated with Roary. A 60% identity threshold was used for defining orthologs in the *Mesorhizobium* pangenome based on a preliminary analysis using various identity thresholds (**Figure S7**). This pangenome of 97 strains was then used as input for a pangenome-wide association study using Scoary, in which the following five strains were marked as isolated from northern sites: the three newly isolated strains (*M*. spp. AR02, AR07, and AR10), as well as *M. loti* 582 and *M. huakuii* 583 that were isolated from *O. kamtschatica* in the Kamchatka Peninsula (Russia) (Safronova et al. 2020). A total of 25 genes were identified as putatively associated with mesorhizobia isolated from northern sites (Benjamini-Hochberg adjusted p-value ≤ 0.01; **Table 3**). Four of the five strains carried at least 24 of these 25 genes, while *M.* sp. AR10 carried only 18, consistent with this strain being the most phylogenetically distant from the others. Notably, using *M.* sp. AR02 as a reference strain, 23 of the 25 genes are found on the symbiotic megaplasmid, with 17 found within 120 kb of the common nodulation genes *nodA*, *nodB*, and *nodC*. Considering that functionally related genes tend to colocalize in bacterial genomes, the genomic location of 23 of the 25 genes putatively associated with northern sites suggests these genes may be related to host interaction or symbiosis, particularly in low temperature environments.

**Table 3.** Genes putatively associated with mesorhizobia isolated from cold climates based on a pangenome-wide association study (Benjamini-Hochberg adjusted p-value ≤ 0.01).

| Locus tag in *M. sp.*<br>AR02 | Replicon in<br>*M. sp.* AR02 | Predicted function | Sensitivity * | Specificity * | *M. huakuii*<br>583 † | *M. loti*<br>582 † | *M. sp.*<br>AR02 † | *M. sp.*<br>AR07 † | *M. sp.*<br>AR10 † |
| --- | --- | --- | --- | --- | --- | --- | --- | --- | --- |
| DBIPINDM_003693 | Chromosome | Nuclear transport factor 2 family protein | 80.0 | 100.0 | + | - | + | + | + |
| DBIPINDM_007274 | pAR02-1 | β-ketoacyl-ACP synthase III | 100.0 | 98.9 | + | + | + | + | + |
| DBIPINDM_007280 | pAR02-1 | Methylmalonyl-CoA mutase subunit beta | 100.0 | 100.0 | + | + | + | + | + |
| DBIPINDM_007281 | pAR02-1 | Sulfate adenylyltransferase subunit NodP | 100.0 | 98.9 | + | + | + | + | + |
| DBIPINDM_007282 | pAR02-1 | Sulfate adenylyltransferase subunit NodQ | 100.0 | 100.0 | + | + | + | + | + |
| DBIPINDM_007283 | pAR02-1 | DUF2061 domain-containing protein | 100.0 | 98.9 | + | + | + | + | + |
| DBIPINDM_007296 | pAR02-1 | Acyltransferase | 100.0 | 98.9 | + | + | + | + | + |
| DBIPINDM_007300 | pAR02-1 | Quinolinate synthase NadA | 100.0 | 96.7 | + | + | + | + | + |
| DBIPINDM_007322 | pAR02-1 | Oxygen-independent coproporphyrinogen III oxidase | 80.0 | 100.0 | + | + | + | + | - |
| DBIPINDM_007351 | pAR02-1 | Indolepyruvate ferredoxin oxidoreductase family protein | 100.0 | 98.9 | + | + | + | + | + |
| DBIPINDM_007357 | pAR02-1 | Serine hydroxymethyltransferase | 100.0 | 100.0 | + | + | + | + | + |
| DBIPINDM_007358 | pAR02-1 | Methionine adenosyltransferase | 100.0 | 97.8 | + | + | + | + | + |
| DBIPINDM_007362 | pAR02-1 | OmpW family protein | 100.0 | 96.7 | + | + | + | + | + |
| DBIPINDM_007376 | pAR02-1 | DNA-binding protein | 100.0 | 96.7 | + | + | + | + | + |
| DBIPINDM_007377 | pAR02-1 | Hypothetical protein | 100.0 | 96.7 | + | + | + | + | + |
| DBIPINDM_007383 | pAR02-1 | Hypothetical protein | 100.0 | 100.0 | + | + | + | + | + |
| DBIPINDM_007752 | pAR02-1 | Hypothetical protein | 80.0 | 100.0 | + | + | + | + | - |
| DBIPINDM_007753 | pAR02-1 | Recombinase family protein | 80.0 | 100.0 | + | + | + | + | - |
| DBIPINDM_007802 | pAR02-1 | Type II toxin-antitoxin system antitoxin | 80.0 | 100.0 | + | + | + | + | - |
| DBIPINDM_007851 | pAR02-1 | OmpT family outer membrane protease | 100.0 | 95.7 | + | + | + | + | + |
| DBIPINDM_007888 | pAR02-1 | Cytochrome c oxidase accessory protein CcoG | 100.0 | 97.8 | + | + | + | + | + |
| DBIPINDM_007933 | pAR02-1 | PIN domain-containing protein | 80.0 | 100.0 | + | + | + | + | - |
| DBIPINDM_007982 | pAR02-1 | TolC family outer membrane protein | 100.0 | 97.8 | + | + | + | + | + |
| DBIPINDM_008005 | pAR02-1 | ATP-binding protein | 80.0 | 100.0 | + | + | + | + | - |
| DBIPINDM_008362 | pAR02-2 | Hypothetical protein | 80.0 | 100.0 | + | + | + | + | - |

\* Sensitivity refers to the percentage of the five strains isolated from arctic or sub-arctic environments that carry the gene. Specificity refers to the percentage of the 92 other strains that lack the gene.
† A positive sign (+) indicates the gene is found in the strain, while a negative sign (-) indicates the gene is not found in the strain.

As discussed above, the primary *nod* locus of all strains is conserved with that of *M*. sp. N33, and thus the Nod factor of all strains is expected to be similar to that of *M*. sp. N33 (Poinsot et al. 2001). Multiple genes were identified as associated with the mesorhizobia isolated from arctic or sub-arctic environments that could be related to Nod factor biosynthesis. These include *nodP* and *nodQ* found within the primary *nod* locus that are required for sulfation of the Nod factor, as well as two genes nearby the primary *nod* locus predicted to encode an acyltransferase and a *β*-ketoacyl-ACP synthase III, which could potentially be involved in the synthesis or addition of the unusual methyl-branched α,β-unsaturated acyl chain of the Nod factor.

Other genes identified by Scoary as associated with mesorhizobia isolated from arctic or sub-arctic environments include those predicted to encode a cytochrome C oxidase accessory protein (CcoG), an inolepyruvate ferredoxin oxidoreductase family protein, and an oxygen-independent coproporphyrinogen III oxidate, all of which could be involved in the functioning of the electron transport chain during nitrogen fixation. We also identified a gene encoding an OmpW family protein; a previous study demonstrated that a *M. japonicum* R7A protein with similarity to OmpW proteins was required for fully effective symbiosis possibly due to a role in protection against oxidative stress (Sullivan et al. 2013). Similarly, a gene predicted to encode a TolC family outer membrane protein was identified, and previous work has shown that a TolC protein is required for symbiosis in *Sinorhizobium meliloti* (Cosme et al. 2008). An omptin family outer membrane protease was also detected, and previous work has shown the up-regulation of an omptin gene in *Bradyrhizobium diazoefficiens* during growth in microoxic conditions (Fernández et al. 2019).

The five strains isolated from northern regions were predicted to encode multiple cold shock proteins, including orthologs of the CspA protein of *Escherichia coli* (Goldstein et al. 1990), and it was previously shown that expression of a cold shock protein is transiently induced in *M.* sp. N33 following cold shock (Ghobakhlou et al. 2015). Although cold shock proteins were not identified as associated with the northern *Mesorhizobium* in the pangenome-wide association study due to these proteins being broadly conserved across the genus, they undoubtedly aid adaptation to low temperatures and northern regions (Prévost et al. 2003).

## CONCLUSIONS

Three *Mesorhizobium* strains (*M.* spp. AR02, AR07, and AR10) from two geographically distinct regions of northern Canada revealed no initially obvious adaptations to growth at low temperatures relative to a common *Mesorhizobium* laboratory strain. However, further studies at lower temperatures (i.e. below 10 C) or for longer durations may reveal psychrotolerant adaptations. A pangenome-wide association study of 97 *Mesorhizobium* strains, of which five were isolated from arctic or sub-arctic climates, identified 25 genes putatively associated with arctic or sub-arctic climates. Of these, 23 are located on the symbiotic megaplasmid and include several genes likely involved in nodulation or nitrogen fixation such as components of the electron transport chain. In future studies, it would be worthwhile to experimentally test the relevance of these genes to SNF in both temperate and low temperature conditions. Of note, we provide evidence that a common set of nodulation genes are highly conserved and shared among the majority of *Mesorhizobium* microsymbionts isolated in arctic and sub-arctic climates, and that these nodulation genes are uncommon in *Mesorhizobium* microsymbionts of other climatic regions. Host plant range appears insufficient by itself to explain the divergence of the nodulation genes of arctic and sub-arctic *Mesorhizobium*. Instead, we hypothesize that this divergence has been driven by climatic factors selecting for increased activity under low temperatures, and that the genes have spread between mesorhizobia across northern environments through horizontal gene transfer events. We anticipate investigating the impact on symbiosis of replacing the *nod* locus of a temperate *Mesorhizobium* strain with the *nod* locus of an arctic *Mesorhizobium* strain, and thus provide an experimental platform for the development of more efficient rhizobial inoculants for cool spring conditions.

## ACKNOWLEDGEMENTS

We thank Robert Colautti for generously assisting with the Nanopore sequencing, as well as Kristy Moniz and Megan Clemens for kindly helping with PCR amplification of the 16S rRNA genes of our *Mesorhizobium* isolates. P.G. is grateful to the Tłıcho Government for permission to conduct research on their lands, and to Karin Clark and the Government of the Northwest Territories’ Environment and Natural Resources division for the use of the Tundra Ecosystem Research Center at Daring Lake.

## CONTRIBUTIONS

G.C.D conceptualized the study. Data collection and analysis was performed by Y.D. and G.C.D. Plant material was collected by V.K.W. and P.G. The first draft of the manuscript was prepared by G.C.D., and all authors contributed to manuscript revision. All authors read and approved the final manuscript.

## DATA AVAILABILITY

Genome assemblies were deposited to the NCBI database under the BioProject accession PRJNA751135. Scripts to repeat the computational analyses, as well as Newick-formatted phylogenies, are available at github.com/diCenzo-Lab/005_2022_northern_mesorhizobia. Bacterial strains are available upon request.

