## Supplemental Tables and Figures for "Whole genome sequencing of three mesorhizobia isolated from northern Canada to identify genomic adaptations promoting nodulation in cold climates"

**This PDF file includes:**

Tables S1 to S3
Figures S1 to S7
Legends for Datasets S1-S11
References for Supplementary Materials

**Other supplementary materials for this manuscript include the following:**

Datasets S1-S11

**Table S1.** Oligonucleotide sequences used in this study.

| Name | Sequence | Reference |
|---|---|---|
| 27f | GAGAGTTTGATCCTGGCTCAG | (Lane 1991) |
| 1495r | CTACGGCTACCTTGTTACGA | (Lane 1991) |
| trnK685F | GTATCGCACTATGTATCATTTGA | (Wojciechowski et al. 2004) |
| trnK2R | CCCGGAACTAGTCGGATGG | (Wojciechowski et al. 2004) |
| matK4La | CCTTCGATACTGGGTGAAAGAT | (Wojciechowski et al. 2004) |

**Table S2.** Carbon sources catabolized by *Mesorhizobium* strains based on Biolog Gen III plates.

| *M*. sp. AR02 | *M*. sp. AR07 | *M*. sp. AR10 |
|---|---|---|
| Common | | |
| α-D-glucose | α-D-glucose | α-D-glucose |
| D-arabitol | D-arabitol | D-arabitol |
| D-fructose | D-fructose | D-fructose |
| D-fucose | D-fucose | D-fucose |
| D-galactose | D-galactose | D-galactose |
| D-maltose | D-maltose | D-maltose |
| D-mannitol | D-mannitol | D-mannitol |
| D-mannose | D-mannose | D-mannose |
| D-sorbitol | D-sorbitol | D-sorbitol |
| D-trehalose | D-trehalose | D-trehalose |
| D-turanose | D-turanose | D-turanose |
| Glycerol | Glycerol | Glycerol |
| L-fucose | L-fucose | L-fucose |
| L-glutamic acid | L-glutamic acid | L-glutamic acid |
| L-lactic acid | L-lactic acid | L-lactic acid |
| L-Malic acid | L-Malic acid | L-Malic acid |
| L-rhamnose | L-rhamnose | L-rhamnose |
| myo-Inositol | myo-Inositol | myo-Inositol |
| N-acetyl-D- galactosamine | N-acetyl-D- galactosamine | N-acetyl-D-galactosamine |
| N-acetyl-D-glucosamine | N-acetyl-D-glucosamine | N-acetyl-D-glucosamine |
| Sucrose | Sucrose | Sucrose |
| Variable | | |
| 3-methyl glucose | α-D-lactose | α-D-lactose |
| Acetic acid | 4-Hydroxy-phenylacetic acid | α-Keto-glutaric acid |
| β-hydroxy-D,L-butyric acid | Acetic acid | β-methyl-D-glucoside |
| D-melibiose | β-methyl-D-glucoside | D-cellobiose |
| D-raffinose | Bromo-succinic acid | D-malic acid |
| Formic acid | D-cellobiose | D-melibiose |
| N-acetyl-B-D-Mannosamine | γ-Amino-butyric acid | Gentiobiose |
| p-Hydroxy-phenylacetic acid | Gentiobiose | Inosine |
| Pectin | Glycyl-L-Proline | L-alanine |
| | Inosine | L-arginine |
| | L-alanine | L-aspartic acid |
| | L-aspartic acid | Methyl pyruvate |
| | L-histidine | Mucic acid |
| | L-serine | N-acetyl-D-mannosamine |
| | Pectin | Tween |

**Table S3.** The *nodA*, *nodB*, and *nodC* genes from the NCBI Nucleotide database that group with the orthologous genes of *M.* sp. AR02, *M.* sp. AR07, and *M.* sp. AR10.

| Accession | Strain* |
|---|---|
| *nodA* | |
| U53327.1 | Rhizobium sp. N33 |
| *nodB* | |
| U53327.1 | *Rhizobium* sp. N33 |
| AF063495.1 | *Rhizobium* sp. CIAM1416 |
| AF063485.1 | *Rhizobium* sp. USDA 4003 |
| AF063482.1 | *Rhizobium* sp. CIAM2032 |
| AF063481.1 | *Rhizobium* sp. CIAM2026 |
| AF063480.1 | *Rhizobium* sp. CIAM0128 |
| AF063479.1 | *Rhizobium* sp. USDA 4007 |
| AF063478.1 | *Rhizobium* sp. USDA 4004 |
| AF063477.1 | *Rhizobium* sp. USDA 3883 |
| AF063476.1 | *Rhizobium* sp. USDA 3876 |
| AF063475.1 | *Rhizobium* sp. USDA 3584 |
| *nodC* | |
| U53327.1 | *Rhizobium* sp. N33 |
| AF063495.1 † | *Rhizobium* sp. CIAM1416 |
| AF063485.1 | *Rhizobium* sp. USDA 4003 |
| AF063482.1 † | *Rhizobium* sp. CIAM2032 |
| AF063481.1 | *Rhizobium* sp. CIAM2026 |
| AF063480.1 | *Rhizobium* sp. CIAM0128 |
| AF063479.1 | *Rhizobium* sp. USDA 4007 |
| AF063478.1 | *Rhizobium* sp. USDA 4004 |
| AF063477.1 | *Rhizobium* sp. USDA 3883 |
| AF063476.1 | *Rhizobium* sp. USDA 3876 |
| AF063475.1 | *Rhizobium* sp. USDA 3584 |
| KJ729212.1 | *Mesorhizobium* sp. CCBAU 75225 |
| KJ729191.1 | *Mesorhizobium* sp. CCBAU 73223 |
| KU729795.1 | *Mesorhizobium* sp. KHD78 |
| KU729794.1 | *Mesorhizobium* sp. KHD77 |
| KU729790.1 | *Mesorhizobium* sp. KHD70 |
| KU729789.1 | *Mesorhizobium* sp. KHD66 |
| KU729788.1 | *Mesorhizobium* sp. KHD65 |
| KU729787.1 | *Mesorhizobium* sp. KHD63 |
| KU729786.1 | *Mesorhizobium* sp. KHD52 |
| KU729785.1 | *Mesorhizobium* sp. KHD34 |

* Strain names are from the NCBI Nucleotide record, and may not reflect current taxonomy.
† Only groups with *M.* sp. AR02, *M.* sp. AR07, and *M.* sp. AR10 when a string of Ns is removed.

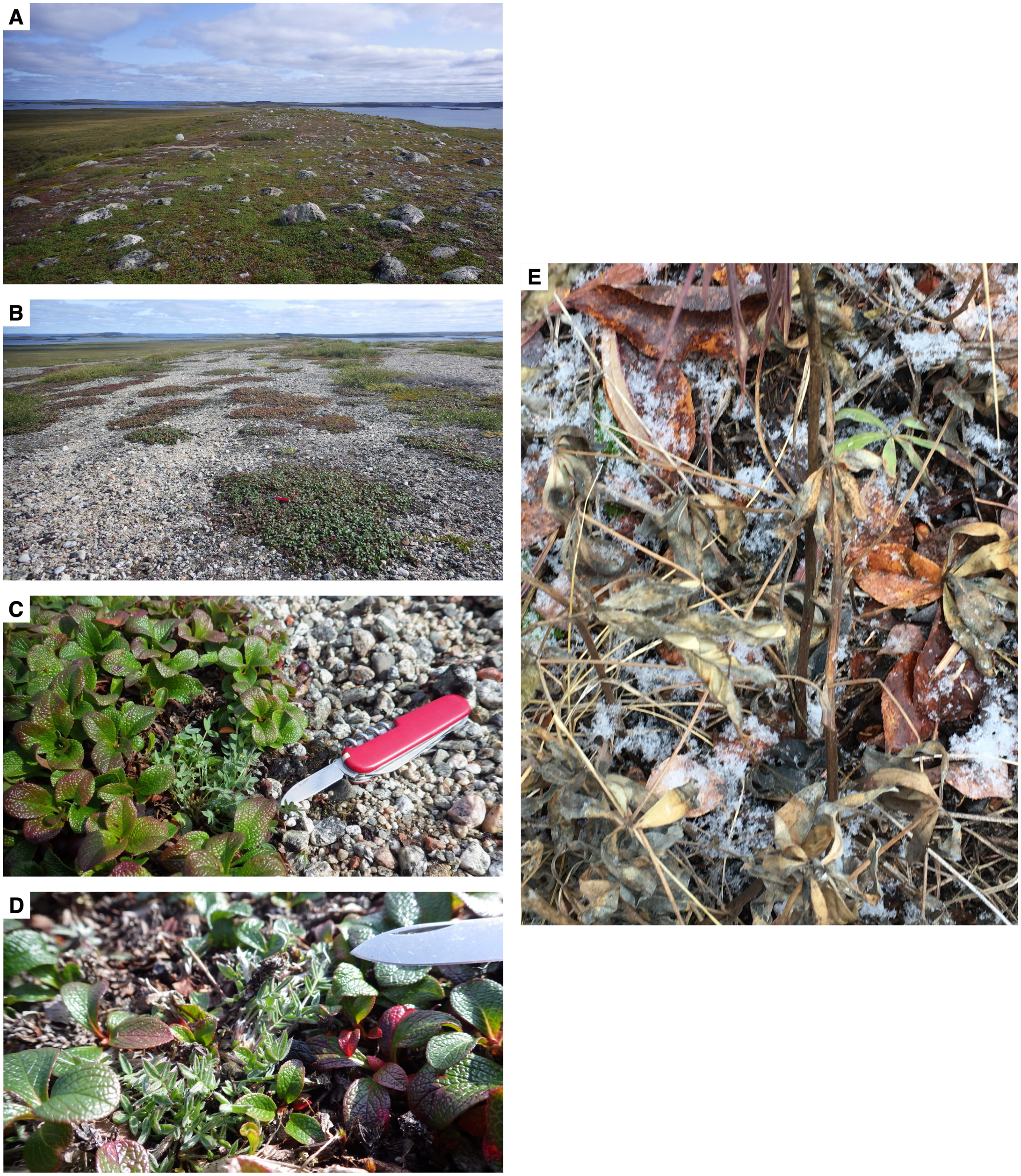

**Figure S1. Photographs from the samples sites.** (**A**,**B**) Photographs of the sampling sites near the Darling Lake Research Station in the Northwest Territories, Canada. (**C**,**D**) Close-up photographs of some of the sampled *Oxytropis nigrescens* plants near the Darling Lake Research Station in the Northwest Territories, Canada. (**E**) Close-up photograp of the sampling site near the Yukon River in Yukon, Canada.

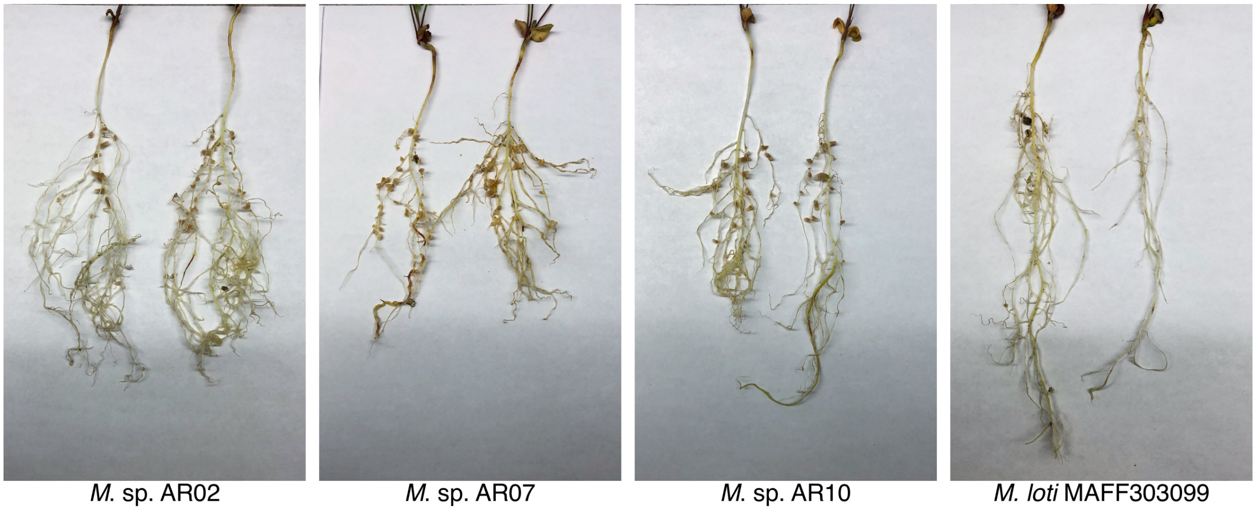


**Figure S2. Sainfoin root systems following inoculation with various *Mesorhizobium* strains.** Pictures of sainfoin (*Onobrychis viciifolia*) plants were taken 36 days following inoculation with *M.* sp. AR02, *M.* sp. AR07, *M.* sp. AR10, or *M. loti* MAFF303099. *M.* sp. AR02, *M.* sp. AR07, and *M.* sp. AR10 were capable of nodulating sainfoin, unlike *M.* loti MAFF303099 that was included as a negative control as a strain whose host range does not include sainfoin.

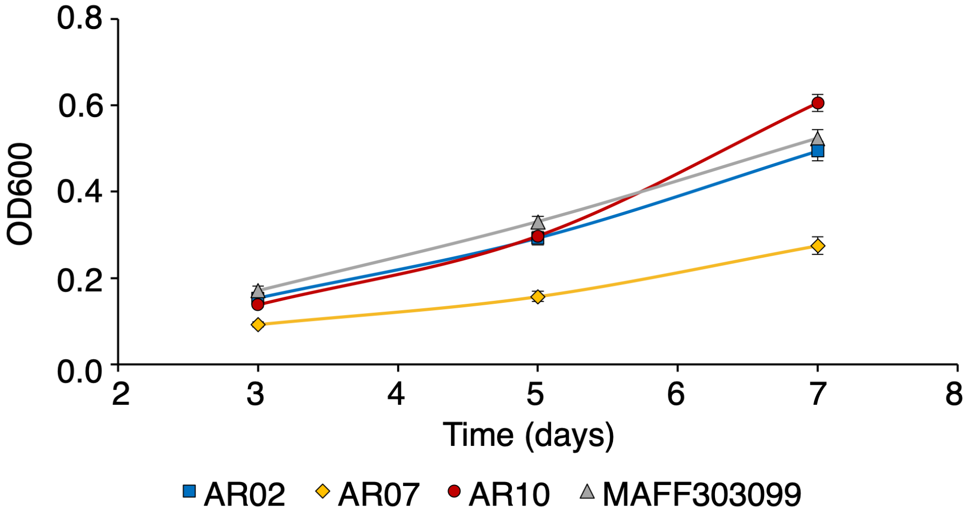


**Figure S3. Growth of *Mesorhizobium* strains at 10˚C.** Cell density ($OD_{600}$) of *Mesorhizobium* cultures following 3, 5, and 7 days of incubation at 10˚C. Starting $OD_{600}$ values were ~ 0.05 for all cultures. Data points represent the average of biological triplicates, with error bars representing the standard deviation. Blue squared – *M.* sp. AR02; yellow diamonds – *M.* sp. AR07; red circles – *M.* sp. AR10; grey triangles – *M. japonicum* MAFF303099.

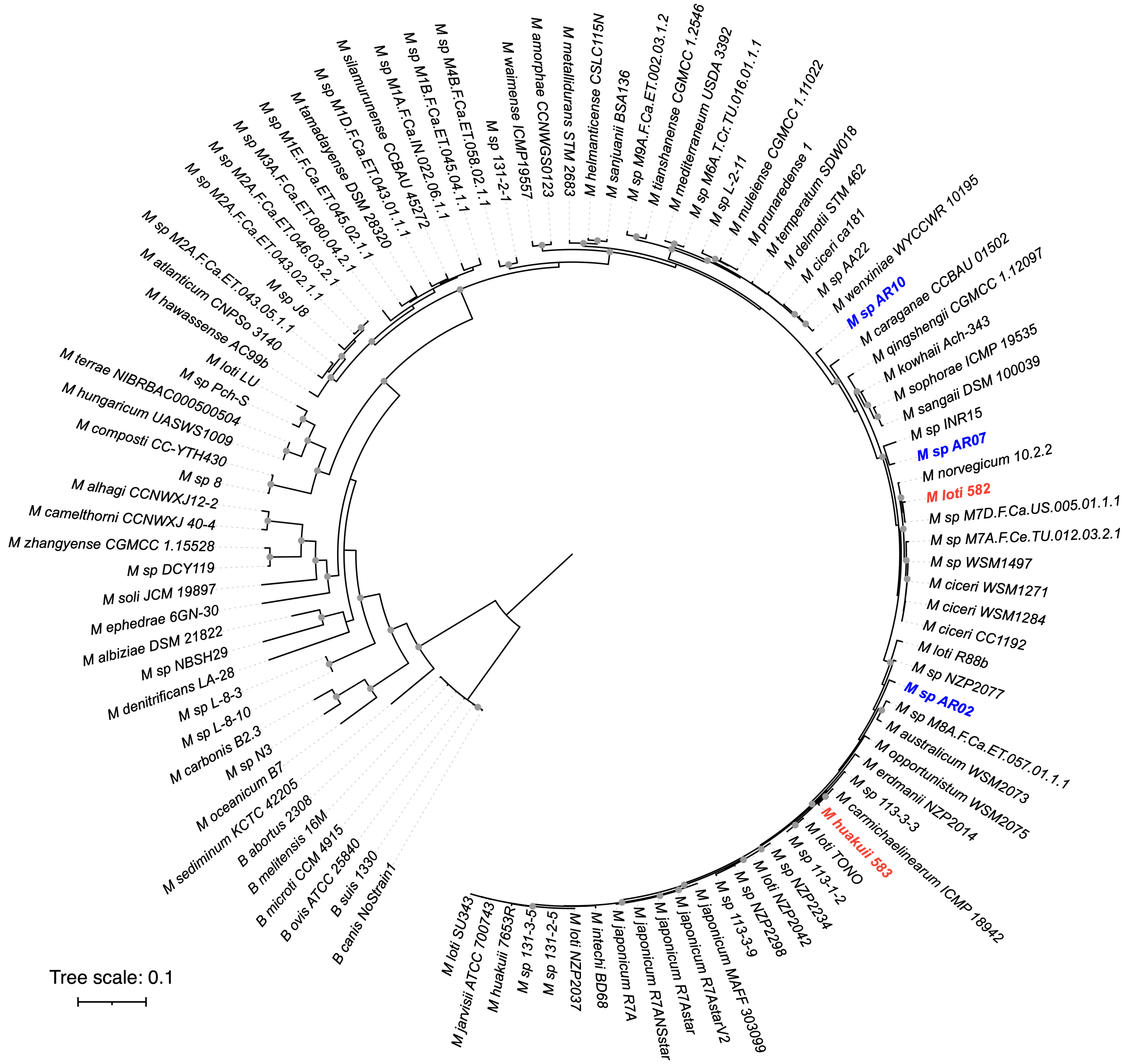


**Figure S4. Rooted phylogeny of the genus *Mesorhizobium*.** A maximum likelihood phylogeny of 97 *Mesorhizobium* strains, rooted with six *Brucella* strains, was prepared from a concatenated alignment of 30 highly conserved proteins (DnaG, Frr, infC, NusA, PyrG, RplA, RplB, RplC, RplD, RplE, RplF, RplK, RplL, RplM, RplN, RplP, RplS, RplT, RpmA, RpoB, RpsB, RpsC, RpsE, RpsI, RpsJ, RpsK, RpsM, RpsS, SmpB, Tsf). Nodes with a boostraap value ≥ 90 are indicated with gray dots. The scale represents the mean number of amino acid substitutions per site. Strains isolated and sequenced in this study are shown in blue, boldface font. *M. loti* 582 and *M. huakuii* 583, also isolated from *Oxytropis* plants in a subarctic climatic region, are shown in red, boldface font.

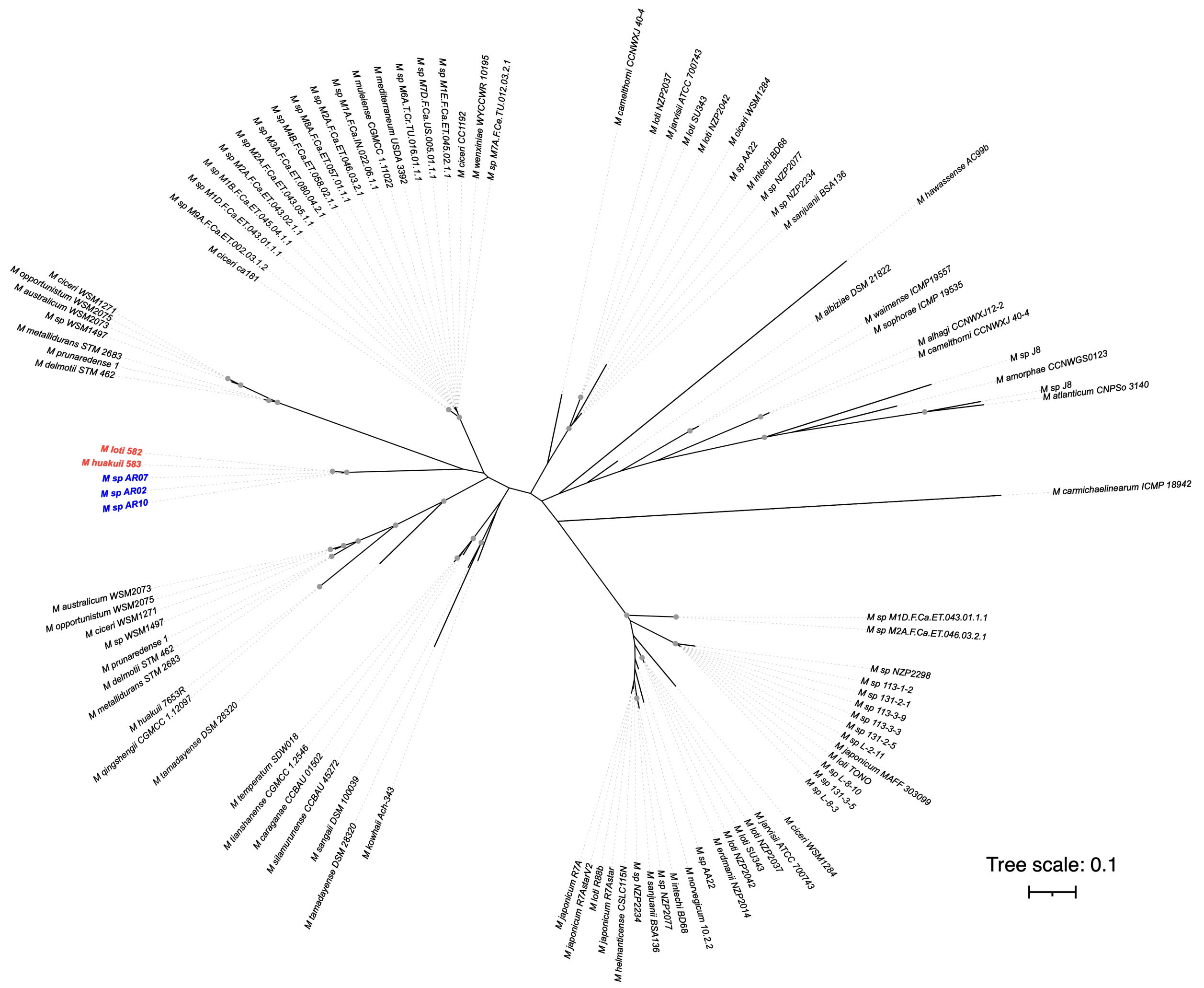

**Figure S5. Phylogeny of the NodA proteins of the mesorhizobia.** A maximum likelihood phylogeny of the NodA proteins of the 97 *Mesorhizobium* strains shown in Figure 2. Nodes with a bootstrap value of ≥ 90 are indicated with gray dots, based on 400 bootstrap replicates. The scale represents the mean number of amino acid substitutions per site. Strains isolated and sequenced in this study are shown in blue, boldface font. *M. loti* 582 and *M. huakuii* 583, also isolated from *Oxytropis* plants in a subarctic climatic region, are shown in red, boldface font.

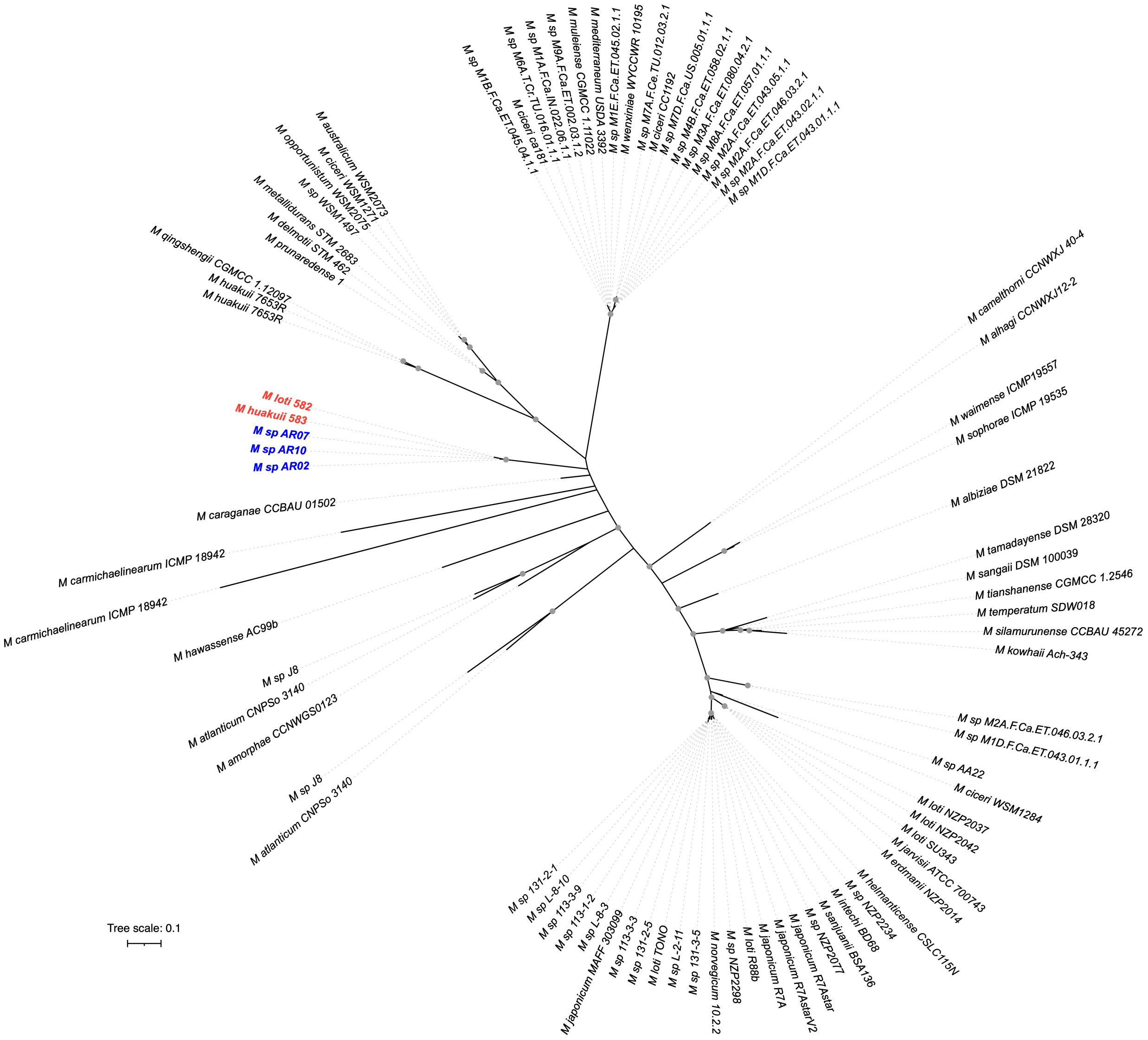


**Figure S6. Phylogeny of the NodB proteins of the mesorhizobia.** A maximum likelihood phylogeny of the NodB proteins of the 97 *Mesorhizobium* strains shown in Figure 2. Nodes with a bootstrap value of ≥ 90 are indicated with gray dots, based on 512 bootstrap replicates. The scale represents the mean number of amino acid substitutions per site. Strains isolated and sequenced in this study are shown in blue, boldface font. *M. loti* 582 and *M. huakuii* 583, also isolated from *Oxytropis* plants in a subarctic climatic region, are shown in red, boldface font.

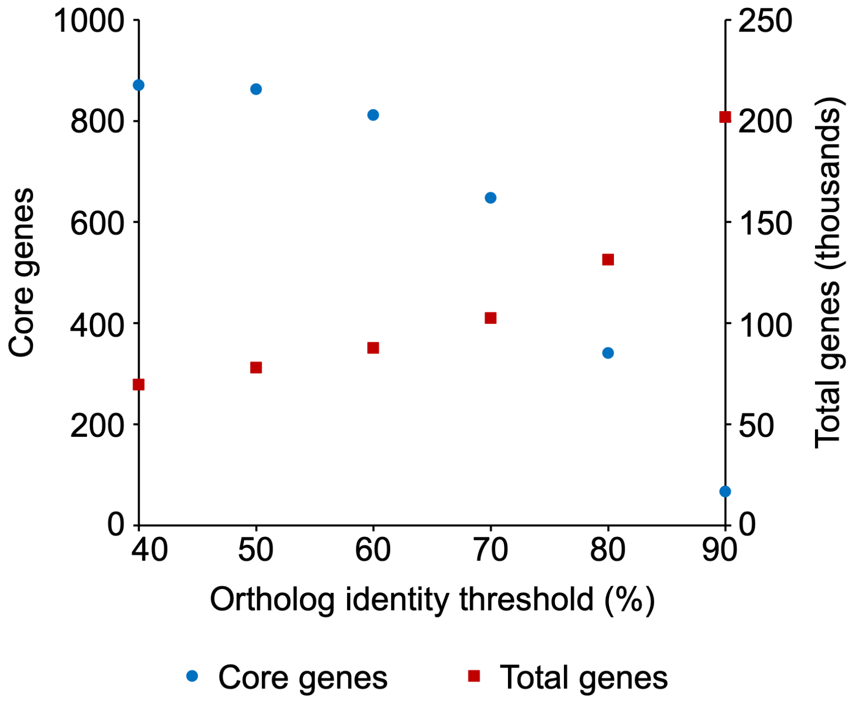


**Figure S7. Impact of ortholog percent identity threshold on pangenome calculation.** Pangenomes of the genus *Mesorhizobium* were calculated using a range of percent identity thresholds for defining orthologs. The number of core genes (blue circles) and total genes (red squares) in the resultant pangenomes are plotted as a function of the ortholog percent identity threshold. An identity threshold of 60% was chosen for downstream analysis as this identity threshold was the highest threshold returning > 90% of core genes identified at an identity threshold of 40%, suggesting this threshold provides a reasonable trade-off between correctly identifying true orthologs and limiting the number of non-orthologous genes being incorrectly grouped as orthologs.

**Dataset S1: Metadata of the *Mesorhizobium* genomes used in the species phylogenetic analysis.** The metadata (as provided by the NCBI Genome database) for the 94 *Mesorhizobium* genomes downloaded from NCBI. All species are named according to their names in the NCBI Genome database at the time of download, which may not necessarily be the currently accepted name.

**Dataset S2: Metadata of the *Brucella* genomes used in the species phylogenetic analysis.** The metadata (as provided by the NCBI Genome database) for the 6 *Brucella* genomes downloaded from NCBI. All species are named according to their names in the NCBI Genome database at the time of download, which may not necessarily be the currently accepted name.

**Dataset S3: NCBI accession numbers for *nodA* genes used in the *nod* gene cluster analysis.** The GenBank and Protein ID accession numbers are provided for the complete set of *nodA* proteins that were downloaded from NCBI for use in the *nod* gene sequence-based cluster analysis.

**Dataset S4: NCBI accession numbers for *nodB* genes used in the *nod* gene cluster analysis.** The GenBank and Protein ID accession numbers are provided for the complete set of *nodB* proteins that were downloaded from NCBI for use in the *nod* gene sequence-based cluster analysis.

**Dataset S5: NCBI accession numbers for *nodC* genes used in the *nod* gene cluster analysis.** The GenBank and Protein ID accession numbers are provided for the complete set of *nodC* proteins that were downloaded from NCBI for use in the *nod* gene sequence-based cluster analysis.

**Dataset S6. Raw output of the Biolog experiment for *M.* sp. AR02.** A Biolog GEN III plate was inoculated with *M.* sp. AR02 and incubated at 27˚C without shaking. Absorbance at 590 nm and 750 nm was measured daily for 10 days. All absorbance readings are provided in this dataset.

**Dataset S7. Raw output of the Biolog experiment for *M.* sp. AR07.** A Biolog GEN III plate was inoculated with *M.* sp. AR07 and incubated at 27˚C without shaking. Absorbance at 590 nm and 750 nm was measured daily for 10 days. All absorbance readings are provided in this dataset.

**Dataset S8. Raw output of the Biolog experiment for *M.* sp. AR10.** A Biolog GEN III plate was inoculated with *M.* sp. AR10 and incubated at 27˚C without shaking. Absorbance at 590 nm and 750 nm was measured daily for 10 days. All absorbance readings are provided in this dataset.

**Dataset S9: Putative prophage regions in *M.* sp. AR02.** Details of the PhiSpy-predicted prophage regions of *M.* sp. AR02 are provided.

**Dataset S10: Putative prophage regions in *M.* sp. AR07.** Details of the PhiSpy-predicted prophage regions of *M.* sp. AR07 are provided.

**Dataset S11: Putative prophage regions in *M.* sp. AR10.** Details of the PhiSpy-predicted prophage regions of *M.* sp. AR10 are provided.
